# Arrestin-3 promotes locomotor sensitization to psychostimulants via JNK signaling in nucleus accumbens

**DOI:** 10.64898/2026.04.28.719936

**Authors:** Mohamed R. Ahmed, Jeffery L. Dunning, Chen Zheng, Sonia Kim, Sebastián Milanés, Christopher Bozorgmehr, Jordan Janzen-Meza, Kathleen Yao, Haoru Li, Vsevolod V. Gurevich, Eugenia V Gurevich

## Abstract

Arrestins play key role in desensitization of G protein-coupled receptors. Direct signaling role of arrestins has also been documented. Two ubiquitously expressed arrestin isoforms, arrestin-2 and -3 (Arr3), perform similarly in receptor desensitization and share many signaling functions, enabling them to substitute for one another. However, certain signaling roles are specific to each isoform. Mice lacking Arr3 (A3KO) show blunted acute responsiveness to the locomotor stimulatory effect of amphetamine (AMPH). Here we demonstrate that AMPH- and cocaine-induced locomotion of A3KO mice is significantly reduced. This loss-of-function phenotype suggests that Arr3-mediated signaling contributes to the effect. Virus-driven expression of Arr3 in caudate-putamen of A3KO and wild type mice suppressed AMPH-induced locomotion. In contrast, restoration of Arr3 in nucleus accumbens rescued locomotor response. Thus, in caudate-putamen Arr3 participates in the desensitization of dopamine receptors, whereas Arr3-dependent signaling in nucleus accumbens underlies the molecular mechanism of the locomotor response and sensitization. Using monofunctional Arr3-derived peptides, we showed that in the nucleus accumbens Arr3 promoted drug-induced locomotor responses via facilitation of JNK3 activation.

---

Arrestins were discovered as negative regulators of G protein-coupled receptors (GPCRs) via homologous desensitization (Carman and Benovic, 1998). In recent years, the role of arrestins as *bona fide* signaling proteins, whose function may or may not depend on their binding to GPCRs, has become appreciated [reviewed in (Gurevich and Gurevich, 2006; DeWire et al., 2007; Gurevich and Gurevich, 2014; Peterson and Luttrell, 2017)]. Many signaling pathways are regulated by GPCRs via G protein- and arrestin-dependent mechanisms, the latter mostly via scaffolding (McDonald et al., 2000; Zhan et al., 2011; Luttrell et al., 2018). The mitogen activated protein (MAP) kinases are the best known arrestin-regulated signaling pathways (Gurevich and Gurevich, 2024). Arrestins facilitate the activation of ERK1/2 (Luttrell et al., 2001; Coffa et al., 2011b; Coffa et al., 2011a; Luttrell and Miller, 2013), JNK (McDonald et al., 2000; Luttrell and Miller, 2013; Zhan et al., 2015; Zhan et al., 2023), and p38 (Chavkin et al., 2014) by scaffolding relevant three-kinase cascades.

Arrestins suppress G protein activation by GPCRs but facilitate different branches of signaling. When the arrestin availability changes, the ultimate outcome depends on the relative contribution of G protein- and arrestin-dependent regulation of a particular pathway, which is cell type-specific (Luttrell et al., 2018). It has been convincingly demonstrated that G protein- and arrestin-dependent signaling respond in the opposite manner to the loss or upregulation of arrestin (Alvarez-Curto et al., 2016; Luttrell et al., 2018). Loss of arrestins results in impaired desensitization and, consequently, enhanced G protein-mediated signaling, whereas overexpression of arrestins has the opposite effect both in cultured cells and living animals (Kohout et al., 2001; Ahn et al., 2003; Bohn et al., 2003; Violin et al., 2008). However, in some cases loss of arrestins results in a loss-of-function phenotype suggesting the involvement of arrestin-dependent signaling rather than the loss of negative regulation (Luttrell et al., 2018).

Being multifunctional regulators of signaling pathways critical for neural adaptations, arrestins are likely involved in the control of responsiveness to acute and long-term effect of psychotropic drugs that directly or indirectly act via GPCRs. Many drugs of different classes produce behavioral sensitization observed as an increased effect of the drug upon repeated administration. That includes dopamine precursor L-DOPA in animal models of Parkinson’s disease (Sgambato-Faure et al., 2005; Cenci, 2007; Ahmed et al., 2010; Ahmed et al., 2015), antipsychotic drugs (Qiao et al., 2012; Li, 2016) and drugs of abuse (Itzhak and Martin, 1999; Vanderschuren and Kalivas, 2000; Valjent et al., 2010; Zurkovsky et al., 2017). We have recently reported that hemiparkinsonian mice lacking arrestin-3 (Arr3) demonstrate reduced responsiveness to dopaminergic stimulation (Ahmed et al., 2024). Arr3 knockout (A3KO) mice also show reduced responsiveness and dampened behavioral sensitization to the psychostimulant drug amphetamine (AMPH) (Beaulieu et al., 2005; Zurkovsky et al., 2017). Phenotypical similarity might be indicative of common underlying mechanisms. Indeed, common neurological background between L-DOPA and antipsychotic-induced sensitization has been reported (Nestler et al., 1999; Graybiel et al., 2000; Schmidt and Beninger, 2006; Savell and Hope, 2023).

Here we explored the signaling mechanisms governing the Arr3-dependent regulation of the locomotor responsiveness to psychostimulants using the locomotor sensitization paradigm with repeated administration of AMPH or cocaine (COC) in mice. This behavioral paradigm is simple, quantitative and allows for the examination of the molecular mechanisms underlying long-term neural plasticity induced by chronic psychostimulant use in living animals. We employed the rescue approach restoring Arr3 to key elements in different parts of the striatal circuitry in the brain of A3KO mice to determine whether it would rescue normal sensitivity to the drugs. We also took advantage of monofunctional Arr3-derived peptides to probe the signaling pathways mediating behavioral effects of psychostimulants. These peptides lack multiple functional capabilities of the full-length Arr3 protein while retaining the ability to modulate the JNK3 activity (Zhan et al., 2016; Perry-Hauser et al., 2022).

Our data demonstrate that the loss-of-function phenotype in A3KO mice is driven by the loss of Arr3 in the nucleus accumbens (NAc), with the largest contribution of the D1 dopamine receptor (D1R)-bearing neurons. We further show that Arr3 promotes locomotor sensitization to both AMPH and COC, similarly to the locomotor sensitization to L-DOPA in hemiparkinsonian mice (Ahmed et al., 2024), via facilitation of JNK3 activation.

## Results

### Arr3 in the striatum dampens psychostimulant-induced hyperlocomotion

Administration of AMPH led to a sharp increase in locomotor activity in rodents. We (Zurkovsky et al., 2017) and others (Beaulieu et al., 2005) previously found that A3KO mice showed blunted locomotor response to AMPH and reduced locomotor sensitization to this drug. Here we sought to investigate the neuronal background of the role of Arr3 in the locomotor effects of psychostimulants.

We employed viral-mediated gene transfer of wild type (WT) Arr3 to the caudate-putamen (CPu) (**Fig. 1A**) to determine whether restoration of Arr3 to striatal neurons would rescue the psychostimulant-induced hyperlocomotion and/or locomotor sensitization (**Fig 1B**). As expected, A3KO mice displayed a blunted locomotor response to AMPH **(Fig. 1C,D,F,G)**. To our surprise, exogenous Arr3 expressed in the CPu of A3KO mice via lentivirus (LV) further reduced already low locomotor response to AMPH (**Fig. 1B,C-F**). The locomotion-suppressing effect of exogenous Arr3 was preserved when the Arr3 expression was confined to D1R-expressing neurons of the CPu by means of the Cre-dependent expression of double-floxed Arr3 driven by adeno-associated virus (AAV) (**Fig. 1B,C,D**). The virus-mediated expression was in both cases localized to the dorsal CPu (**Fig. 1B,F-K**). The AAV-mediated expression was confined to Cre-positive neurons (**Fig. 1I,K and S1C-E**). The expression levels of the Arr3 driven by LV under CMV promoter, which ensured pan-neuronal expression, and AAV driving Cre-dependent expression selective for D1R-bearing striatal neurons were comparable: ∼50% of the endogenous level of Arr3 for LV and ∼80% for the AAV (**Fig. 1L**).

**Figure 1.**
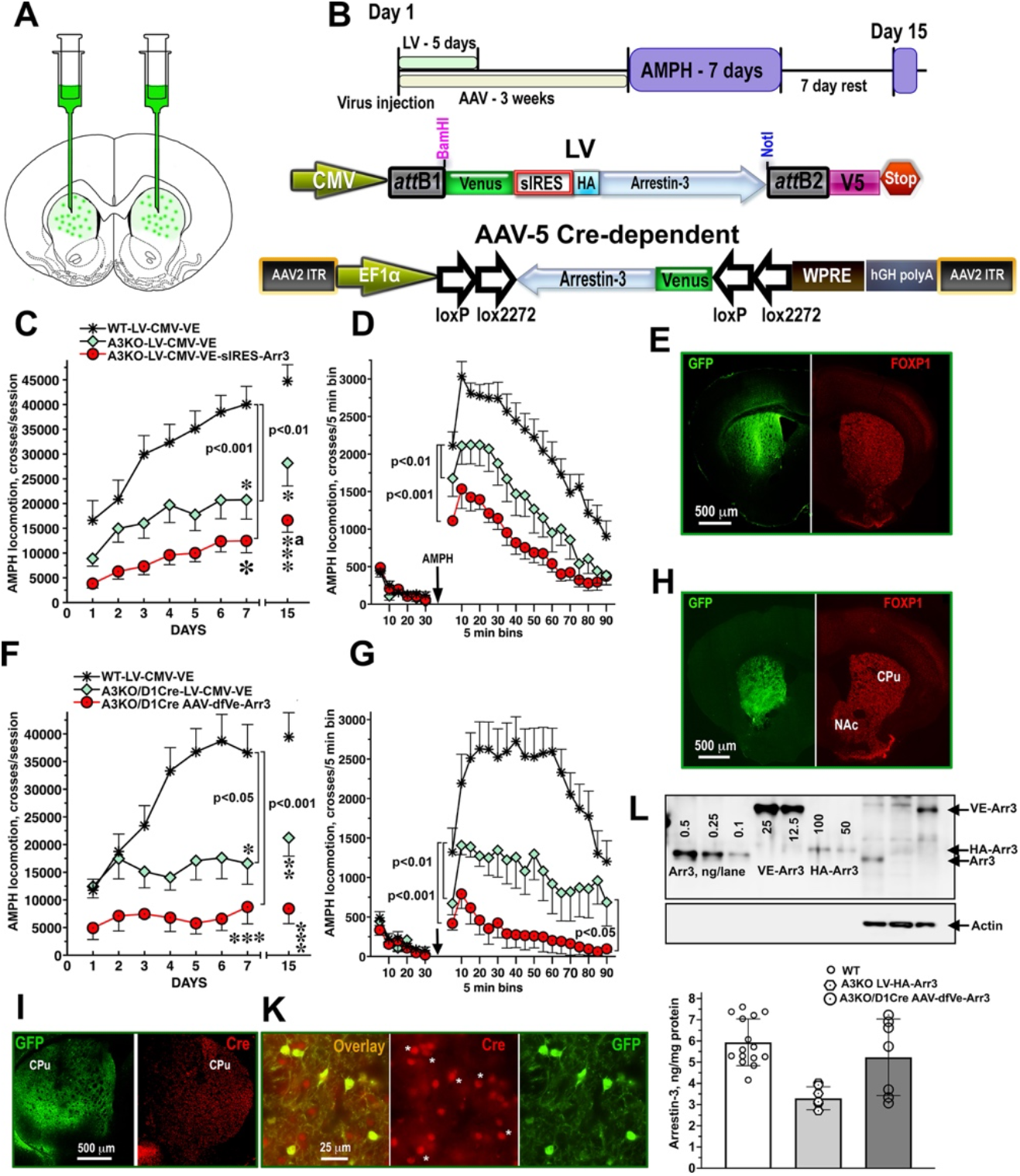
Arrestin-3 in the CPu of A3KO mice further suppresses blunted AMPH-induced locomotion. (**A**) A3KO mice were injected bilaterally into the CPu with the LV or AAV encoding Venus (VE) or wild type (WT) Arr3. WT mice received VE. **(B)** Experimental time course (top) and the viruses used. LV expressed under control of the CMV promoter Venus and co-cistronically Arr3-HA preceded by superIRES (Bochkov and Palmenberg, 2006; Ahmed et al., 2024). The AAV was designed for Cre-dependent expression (double-floxed open reading frame-inverted) of VE-Arr3 under control of the EF1α promoter. **(C)** LV-injected mice were treated with AMPH (3 mg/kg, daily for 7 days). Total AMPH-induced locomotion was measured for 90 min after administration of AMPH. Mice were challenged with AMPH following a 7-day withdrawal on Day 15. (**D**) The basal and AMPH-induced locomotor activity on Day 6 for the entire testing period of 30 min (basal activity) and 90 min (AMPH-induced activity) in A3KO and WT mice shown in **C**. **(E)** A photomicrograph showing the LV-driven Arr3 expression (detected by co-cistronic Venus, green). The sections were co-stained for a marker of medium spiny neurons FOXP1 (red). **(F)** Total AMPH-induced locomotion across days in WT and A3KO mice and A3KO mice injected with the AAV. (**G**) The basal and AMPH-induced locomotor activity on Day 6 in A3KO and WT groups shown in **F**. **(H)** A photomicrograph showing the AAV-driven VE-Arr3 expression (labeled for GFP, green) in striatal neurons stained for FOXP1 (red). **(I**,**K**) Co-expression of Venus with Cre in CPu. Low (**I**) and high (**K**) magnification photomicrographs of a section co-stained for Venus and Cre are shown. Asterisks in **K** indicated Cre-positive cells expressing Venus. **(L)** The level of LV- and AAV-driven expression of Arr3 measured by Western blot in comparison with the endogenous Arr3 in CPu (N=5-12). The behavioral data were analyzed by two-way repeated measure ANOVA with Group (Genotype + injected virus) as between group and Day or Bin as within group factor followed by Bonferroni’s post hoc comparisons for the effects of Group. Significance values next to brackets refer to the differences between Groups across Days or Bins. For the Challenge Day (Day 15), **\*** - p<0.05, **\*\***- p<0.01, **\*\*\*\*** - p<0.001 to the respective WT group; **a** - p<0.05 to A3KO-VE.

Next, we tested the effect of Arr3 overexpression in WT mice with normal complement of endogenous Arr3 on psychostimulant-induced hyperlocomotion. We found that LV-mediated expression of Arr3 in all CPu neurons caused a modest transient reduction in AMPH-Induced locomotion (**Fig. 2A,B**). The Cre-dependent expression of Arr3 in D1R-bearing neurons in the CPu resulted in a significant reduction in the locomotor response to AMPH and almost completely suppressed sensitization (**Fig. 2C,D**). In contrast, expression of Arr3 in the CPu neurons expressing Cre under control of the adenosine A2A receptor promotor in the D2 dopamine receptor (D2R)-bearing neurons was entirely ineffective (**Fig. 2E,F**). Interestingly, the level of Arr3 in D1R Cre-dependent expression was substantially lower than that of the A2A-dependent expression **(Fig. 2G,H)**. Thus, in the CPu Arr3 functions to limit the locomotor response to psychostimulants acting in the direct pathway D1R-bearing neurons.

**Figure 2.**
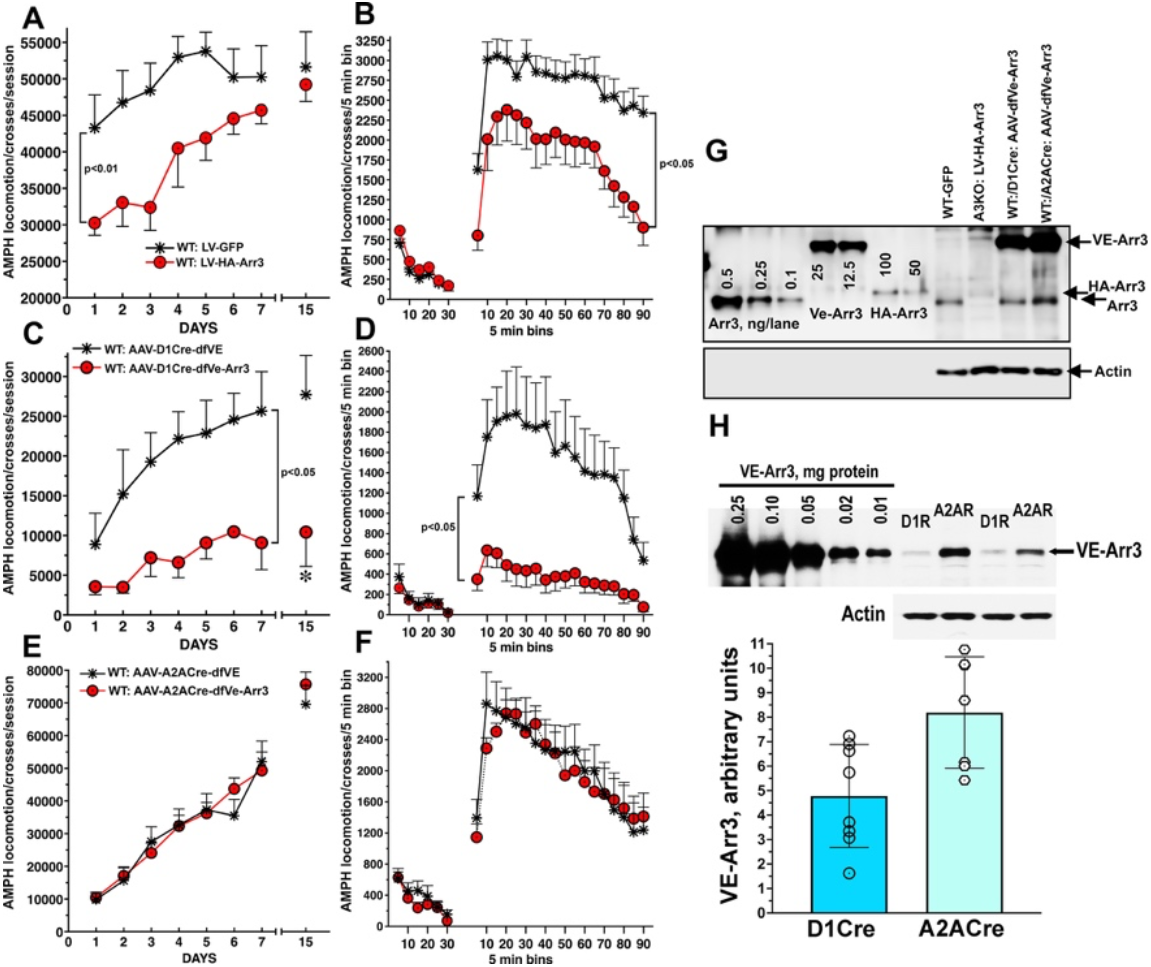
Arr3 in the D1R-bearing neurons in the CPu suppresses the locomotor effect of psychostimulants. (**A**,**B**) WT mice were injected bilaterally into the CPu as with LV encoding VE or VE-Arr3. (**A**) Total AMPH-induced locomotion measured for 90 min after administration of the drug. (**B**) The basal and AMPH-induced locomotor activity on Day 3 for the entire testing period of 30 min (basal activity) and 90 min (AMPH-induced activity) in the groups shown in **A**. The data were analyzed by two-way repeated measure ANOVA with Group (Genotype + injected virus) as between group and Day or Bin as within group factor. Significance values next to brackets refer to the F value differences between Groups across Days or Bins. * - p<0.05 according to two-tailed Student’s t-test for the Challenge Day (Day 15). (**C**,**D**) WT mice hemizygous for Cre under D1R promotor (D1R-Cre) were injected into the CPu with adenoviruses encoding double-floxed open reading frame-inverted Venus (dfVE) or WT Arr3 (dfArr3). (**C**) Total AMPH-induced locomotion measured for 90 min after administration of the drug. Statistical analysis was the same as in **A**. (**D**) The basal and AMPH-induced locomotor activity on Day 4 for the entire testing period of 30 min (basal activity) and 90 min (AMPH-induced activity) in the groups shown in **C**. Statistical analysis was the same as in **B**. (**E**,**F**) WT mice hemizygous for Cre under A2A adenosine receptor promotor (A2A-Cre) were injected into the CPu dfVE or dfA3WT). (**E**) Total AMPH-induced locomotion measured for 90 min after administration of the drug. Statistical analysis was the same as in **A**. (**F**) The basal and AMPH-induced locomotor activity on Day 5 for the entire testing period of 30 min (basal activity) and 90 min (AMPH-induced activity) in the groups shown in **E**. (**G**) The expression of Arr3 in the CPu of WT mice following injection of LV-HA-Arr3 of Cre-dependent AAV-VE-Arr3 detected with anti-Arr3 antibody. Representative Western blot includes the standard samples to indicate the position ofthe bands: purified Arr3, ng/lane; VE-Arr3 or HA-Arr3 expressed in HEK DKO cells. (**H**) The expression of exogenous VE-Arr3 driven by Cre-dependent AAV in D1R-Cre and A2AR-Cre mice detected with anti-GFP antibody. Upper panel: representative Western blot; lower panel – quantification of the Western blot data (N=7).

### Arr3 in the nucleus accumbens is required for psychostimulant-induced hyperlocomotion

Loss-of-function phenotype of A3KO mice suggests that Arr3 is necessary for the full-scale psychostimulant-induced hyperlocomotion and sensitization. Since the expression of Arr3 in the CPu further reduced the locomotor effects, the loss of Arr3 in the CPu was not the cause of phenotype. We next tested whether restoration of Arr3 selectively to the nucleus accumbens (NAc) would rescue the response. We found that LV-driven expression of Arr3 in NAc (**Fig. 3A,B**) of A3KO mice significantly elevated their locomotor responsiveness to AMPH, albeit not to the level of that of WT mice (**Fig. 3C,D**). Interestingly, contrary to the previous report (Bohn et al., 2003), we found that A3KO mice showed minimal hyperlocomotion and sensitization in response to COC (**Fig. 3E,F**). LV-mediated expression of Arr3 in NAc also fully rescued the locomotor response and sensitization to COC (**Fig. 3E,F**).

**Figure 3.**
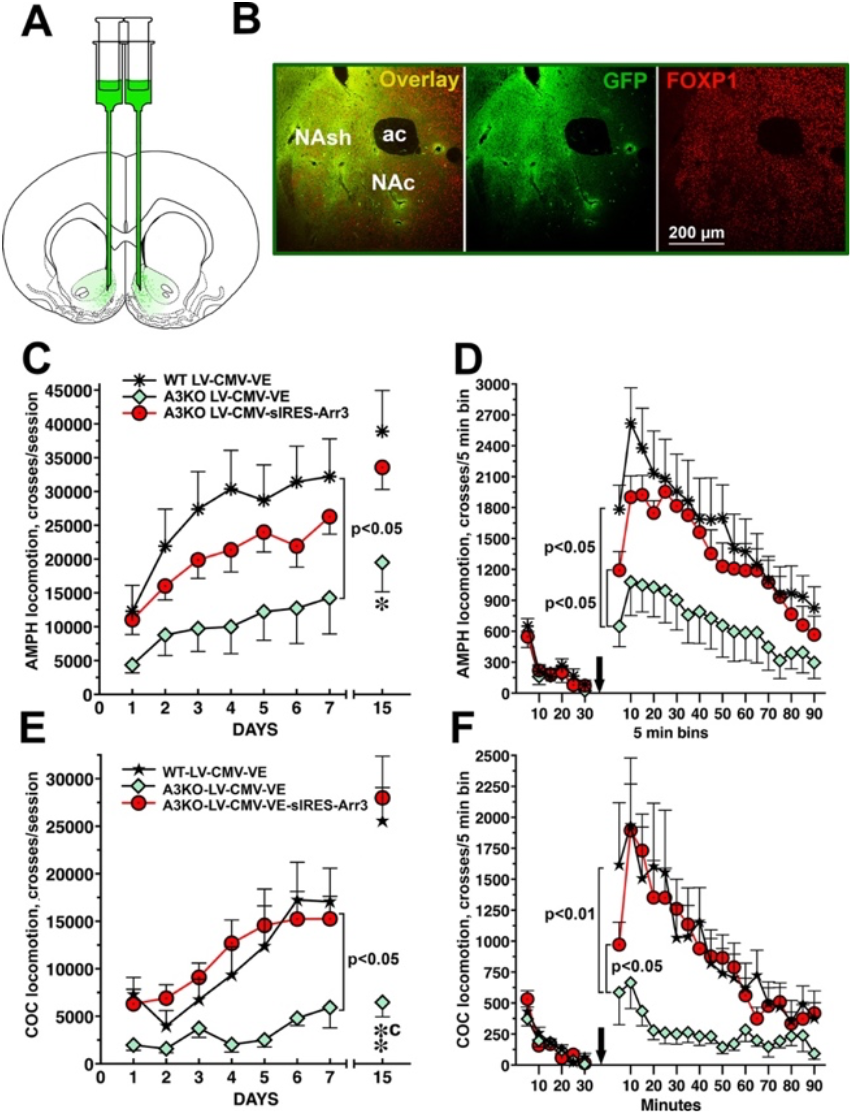
Restoration of Arr3 to the NAc rescues psychostimulant-induced hyperlocomotion. **(A)** Mice were injected bilaterally into the NAc targeting primarily the shell. **(B)** LV-mediated expression of Venus in NAc detected by immunohistochemistry for Venus (green). FOXP1 was used as a marker for medium spiny neurons (red). **(C**,**D)** A3KO mice were injected bilaterally into NAc with the LV encoding either VE or VE-Arr3. WT mice received VE. Mice were treated with amphetamine (AMPH) (3 mg/kg, daily for 7 days and challenged with the same dose on day 15). **(C)** Total AMPH-induced locomotion measured for 90 min after administration of the drug for each testing day and the Challenge Day. (**D)** The basal and AMPH-induced locomotor on Day 5 for the entire testing period of 30 min (basal activity) and 90 min (AMPH-induced activity) in A3KO and WT mice shown in **D**. Arrow indicates the time of AMPH administration. **(E**,**F**) A3KO mice were injected bilaterally into NAc with the LV encoding VE or VE-Arr3. WT mice received VE. Mice were treated with cocaine (COC) (20 mg/kg, daily for 7 days and challenged with the same dose on day 15). **(E)** Total COC-induced locomotion measured for 90 min after administration of the drug for each testing day and the Challenge Day. (**F**) The basal and COC-induced locomotion on Day 5 for the entire testing period of 30 min (basal activity) and 90 min (COC-induced activity) in A3KO and WT mice shown in **F**. Arrow indicates the time of COC administration. The data were analyzed by two-way repeated measure ANOVA with Group (Genotype + injected virus) as between group and Day or Bin as within group factor followed by Bonferroni’s post hoc comparisons for the effects of Group. Significance values next to brackets refer to the differences between Groups across Days or Bins. For the Challenge Day (Day 15), * - p<0.05, **- p<0.01, to the respective WT group; c - p<0.001 to A3KO-VE.

To further explore the neuronal basis of Arr3 effects, we constructed LVs encoding Arr3 under control of dynorphin (DYN) or enkephalin (ENK) promoters to achieve neuron-specific expression (**Fig. 4A**). We previously demonstrated that these promoters drive cell-specific expression in neurons expressing DYN or ENK (Ahmed et al., 2024). We found that the expression of Arr3 in DYN-expressing neurons not only afforded full behavioral rescue but lead to a stronger locomotor response to AMPH than in WT control (**Fig. 4B,C**). In contrast, the Arr3 expression in ENK-expressing neurons resulted in only partial rescue (**Fig. 4B,C**). The levels of Arr3 expression driven by DYN and ENK promotors were similar (**Fig. 4 F,G**).

**Figure 4.**
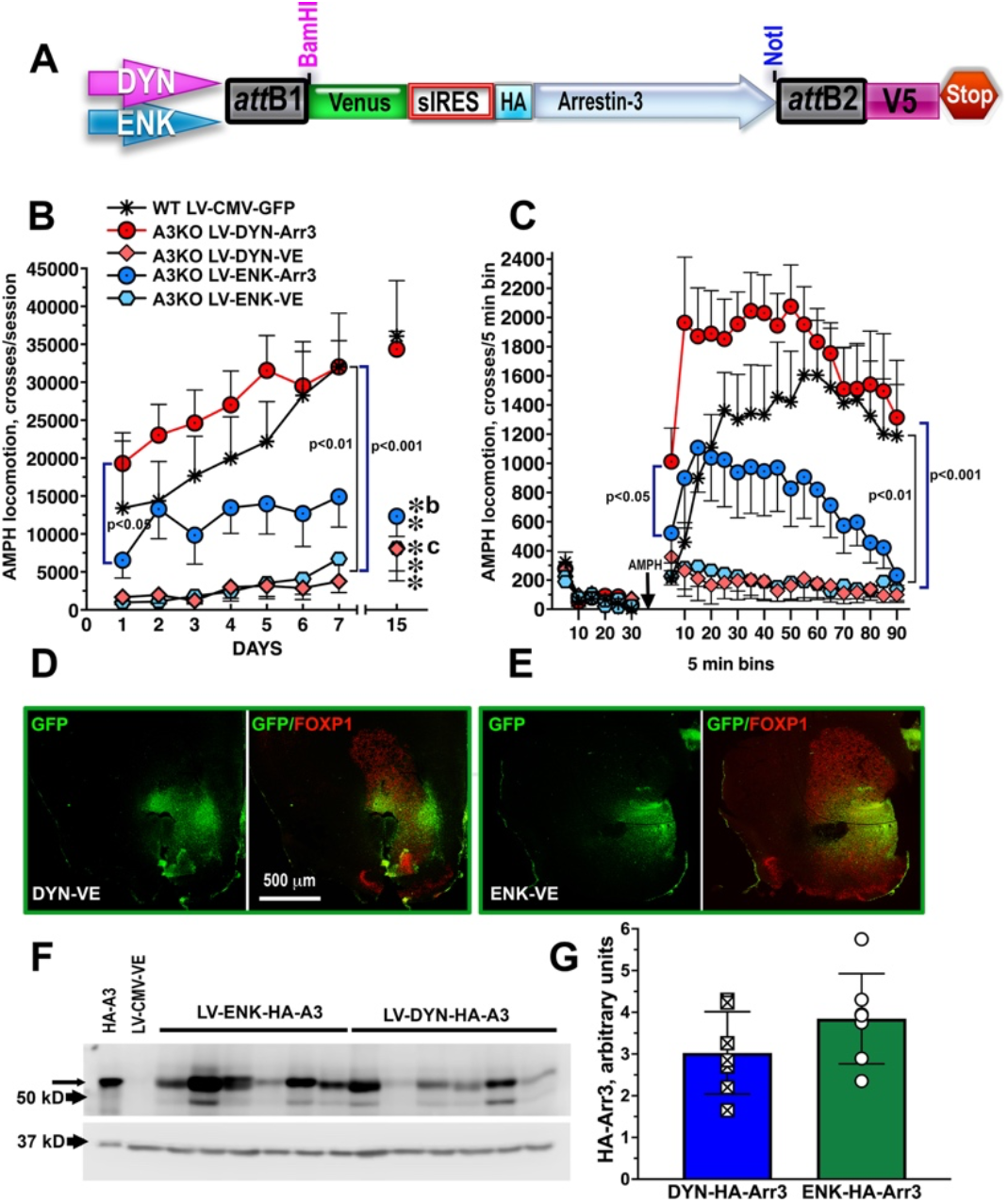
Arr3 in the dynorphine-expressing neurons in NAc promotes the locomotor effect of amphetamine. **(A)** The design of LVs encoding HA-Arr3 with co-cistronic expression of Venus. **(B**,**C)** A3KO mice were injected bilaterally into NAc with LV encoding Venus (VE) or VE+HA-Arr3 under control of dynorphin (DYN) or enkephalin (ENK) promotor. The AMPH (3 mg/kg)-induced locomotion was measured for 7 days. **(B)** Total AMPH-induced locomotion measured for 90 min after administration of the drug. Mice were challenged with AMPH following a 7-day withdrawal on Day 15. (**C**) The basal and AMPH-induced locomotion on Day 5 for the entire testing period of 30 min (basal activity) and 90 min (AMPH-induced activity) in the mice shown in **B. \*\*** - p<0.01, **\*\*\*** - p<0.001 to WT-LV-CMV-VE; b – p<0.01, **c** – p<0.001 A3KO-LV-DYN-Arr3 to A3KO-LV-DYN-VE and A3KO-LV-ENK-VE. **(D**,**E)** Photomicrographs showing the expression of HA-Arr3 in the NAc of D1Cre **(D)** and A2ACre **(E)** mice. The sections were stained to detect Venus co-cistronically expressed with Arr3 (green) and co-stained for FOXP1, a marker of striatal neurons. **(F**,**G)** Detection of HA-Arr3 expression driven by DYN or ENK promoters by Western blot with anti-HA antibody. **(F)** Representative Western blot. **(G)** Quantification of the Western blot data (N=6-7).

### Arr3 in D1R-bearing neurons in NAc is essential for the locomotor effects of psychostimulants

The neuronal composition of the NAc is more complex than that of the CPu, and there is a substantial overlap in the expression of the dopamine receptor subtypes and neuropeptides (Lu et al., 1998; Zhou et al., 2003; Kupchik et al., 2015). To further test the neuronal basis of the Arr3 function, we compared the role of Arr3 in neuronal subtypes of the NAc in the locomotor effects of psychostimulant drugs. The AAV construct with double-floxed open reading frame-inverted design (**Fig. S1A**) ensured strictly Cre-dependent expression (**Fig. S1B-F**).

The AAV-mediated expression of Arr3 in the D1R-bearing neurons in NAc fully rescued the hyperlocomotor response to AMPH (**Fig. 5A,B**), whereas its expression in the D2R-bearing neurons was ineffective (**Fig. 5C,D**). Arr3 was selectively expressed in NAc (**Fig. 5E**) in D1R-Cre mice and in A2A-Cre mice (**Fig. 5 F**) at comparable levels (**Fig. 5 F**).

**Figure 5.**
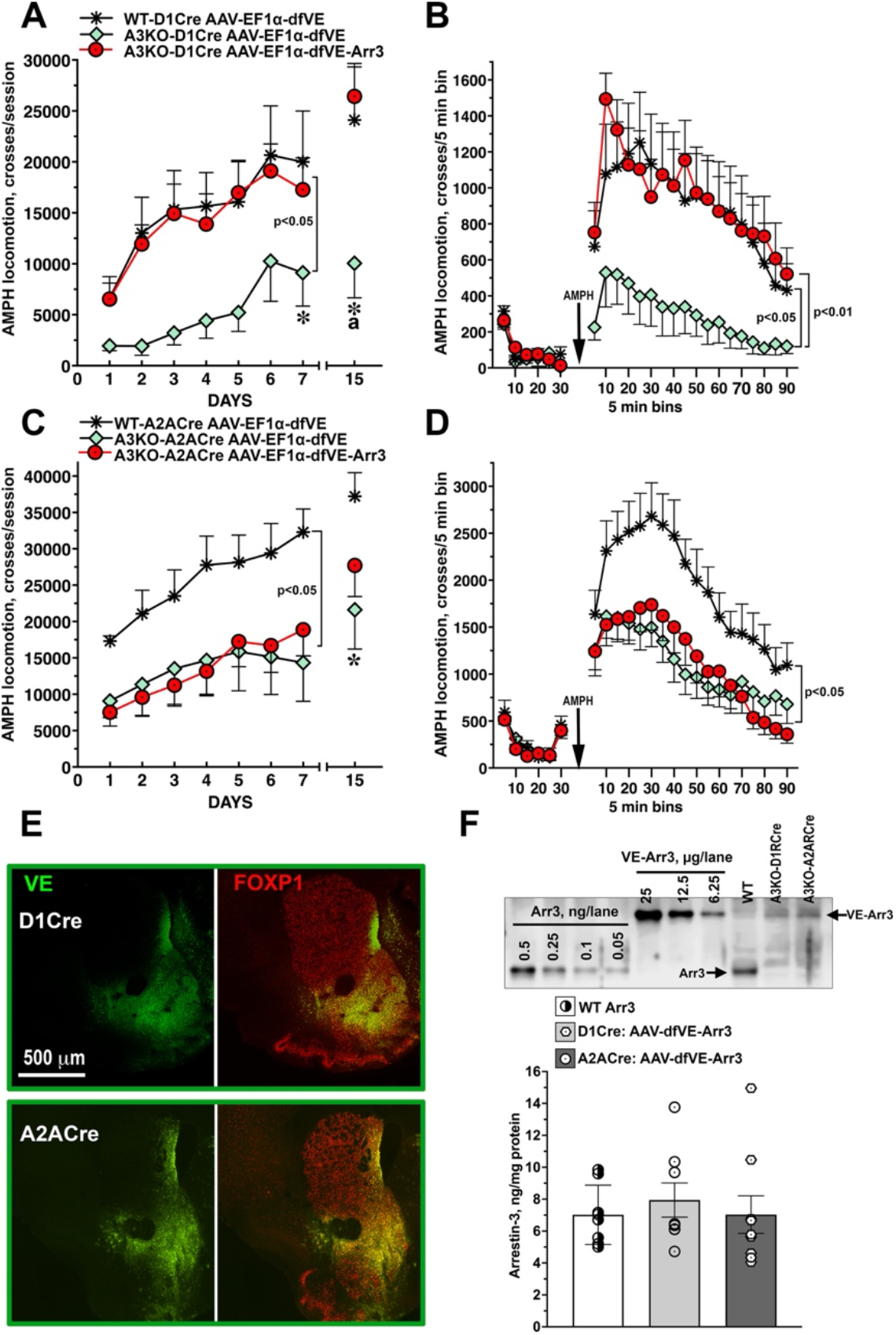
Arr3 in the D1R-bearing neurons in NAc is required for the the locomotor effect of amphetamine. (**A**,**B**) A3KO mice hemizygous for Cre under D1R promotor (D1R-Cre) were injected into the nucleus accumbens (NAc) with AAVs encoding double-floxed open reading frame-inverted Venus (dfVE) or WT Arr3 (dfArr3). (**A**) Total AMPH-induced locomotion measured for 90 min after administration of the drug. (**B**) The basal and AMPH-induced locomotor activity on Day 5 for the entire testing period of 30 min (basal activity) and 90 min (AMPH-induced activity) in the mice shown in **A**. The data were analyzed by two-way repeated measure ANOVA with Group (Genotype + injected virus) as between group and Day or Bin as within group factor. Significance values next to brackets refer to the differences between Groups across Days or Bins. The p value in **A** applies to both WT-VE -A3KO-VE and A3KO-Arr3 - A3KO-VE. * - p<0.05 to WT-VE, a - p<0.05 to A3KO-Arr3 according to Bonferroni’s post-hoc test for the Challenge Day (Day 15). (**C**,**D**) A3KO mice hemizygous for Cre under A2A adenosine receptor promotor (A2A-Cre) were injected into NAc with dfVE or dfArr3. (**C**) Total AMPH-induced locomotion measured for 90 min after administration of the drug. Statistical analysis was the same as in **A**. (**D**) The basal and AMPH-induced locomotor activity on Day 5 for the entire testing period of 30 min (basal activity) and 90 min (AMPH-induced activity) in the mice shown in **C**. Statistical analysis was the same as in **A, B**. In **C** and **D**, the p values apply to the differences A3KO-VE – WT-VE and A3KO-Arr3 - WT-VE. * - p<0.05 to WT-VE. (**E**) –The expression of exogenous VE-Arr3 in NAc driven by D1R-Cre and A2AR-Cre detected by immunohistochemistry. The sections were co-stained for the marker of MSNs FOXP1 (red). (**F**) The level of Cre driven expression of VE-Arr3 in comparison with the endogenous Arr3 measured by Western blot with anti-Arr3 antibody. Upper panel: representative western blot; lower panel – quantification of the Western blot data (N=8-10).

The AAV-mediated expression of Arr3 in the D1R-bearing neurons in NAc also rescued the hyperlocomotor response to another psychostimulant drug COC (**Fig. 6A,B**), while the expression of Arr3 in the D2R-bearing neurons had no effect (**Fig. 6C,D**).

**Figure 6.**
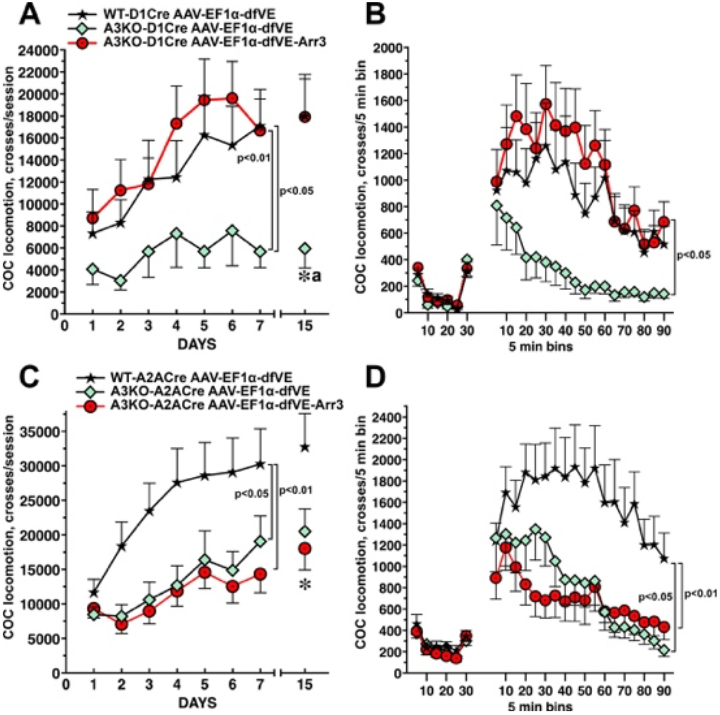
Arr3 in the D1R-bearing neurons in NAc is required for the locomotor effect of cocaine. (**A**,**B**) A3KO mice hemizygous for Cre under D1R promotor (D1R-Cre) were injected into NAc with AAVs encoding double-floxed open reading frame-inverted Venus (dfVE) or WT Arr3 (dfArr3). (**A**) Total COC-induced locomotion measured for 90 min after administration of the drug. (**B**) The basal and COC-induced locomotor activity on Day 5 for the entire testing period of 30 min (basal activity) and 90 min (AMPH-induced activity) in mice shown in **A**. The data were analyzed by two-way repeated measure ANOVA with Group (Genotype + injected virus) as between group and Day or Bin as within group factor. Significance values next to brackets refer to the differences between Groups across Days or Bins. * - p<0.05 to WT-VE, a - p<0.05 to A3KO-Arr3 according to Bonferroni’s post-hoc test for the Challenge Day (Day 15). (**C**,**D**) A3KO mice hemizygous for Cre under A2A adenosine receptor promotor (A2A-Cre) were injected into NAc with dfVE or dfArr3 AAVs. (**C**) Total COC-induced locomotion measured for 90 min after administration of the drug. Statistical analysis was the same as in **A**. (**D**) The basal and COC-induced locomotor activity on Day 5 for the entire testing period of 30 min (basal activity) and 90 min (COC-induced activity) in the mice shown in **C**. Statistical analysis was the same as in **A, B**. * - p<0.05 to WT-VE.

### Arr3 mediates locomotor effects of psychostimulants via JNK activation

The reduced responsiveness of A3KO mice to psychostimulants bears a strong resemblance to diminished responsiveness and sensitization to L-DOPA in hemiparkinsonian A3KO mice (Ahmed et al., 2024). We found that Arr3 supports L-DOPA-induced behaviors by facilitating JNK3 activation (Ahmed et al., 2024). Therefore, we tested whether the Arr3-dependent JNK3 activation is involved in the psychostimulant-induced hyperlocomotion and sensitization to psychostimulants. To this end, we used a tool we have developed, a 25-residue Arr3-derived peptide T1A (**Fig. S2A**) that facilitates JNK activation like full-length Arr3 (**Fig. S2B**,**C**) (Zhan et al., 2016; Perry-Hauser et al., 2022; Ahmed et al., 2024). We used an AAV for pan-neuronal expression (**Fig. 7A)** of Venus-tagged full-length Arr3, or Venus-tagged T1A (VE-T1A). A homologous peptide derived from arrestin-2, VE-B1A, unable to activate JNK (Zhan et al., 2016) or rescue JNK-dependent behavior (Ahmed et al., 2024) was used as a negative control. The advantage of T1A is that it is a monofunctional tool that facilitates JNK3 activation (**Fig. S2B**,**C**) but does not perform multiple other functions of the full-length Arr3 (**Fig. S3A-F**).

**Figure 7.**
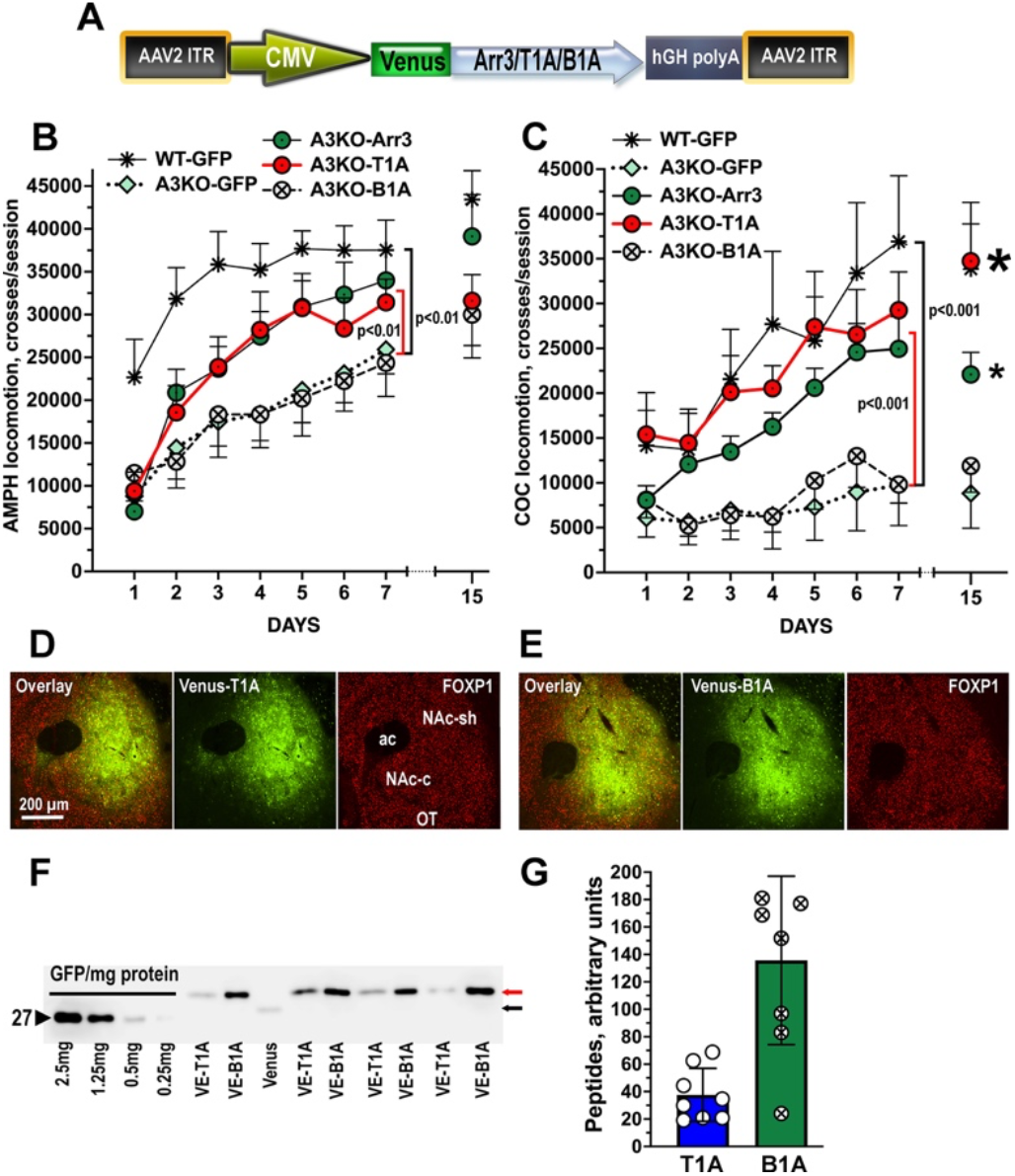
Arrestin-3-derived JNK-activating peptide T1A restores the locomotor effects of psychostimulants. **(A)** Design of AAV-5 for pan-neuronal expression of VE-Arr3, VE-T1A and VE-B1A under control of the CMV promoter. **(B**,**C**) A3KO mice were injected with AAV encoding VE, VE-Arr3, VE-T1A, or VE-B1A and tested for drug-induced locomotion for 7 consecutive days and then for 1 day after a 7-day break. (**B**) AMPH-induced locomotion. Black bracket shows the significant difference between WT and A3KO-VE and A3KO-VE-B1A groups; red bracket – between A3KO-VE-Arr3 or A3KO-VE-T1A and A3KO-VE or A3KO-VE-B1A controls. (**C**) COC-induced locomotion. Long bracket shows significant difference between the A3KO-VE or A3KO-VE-B1A group. Shorter bracket shows significant difference between both A3KO-VE-Arr3 and A3KO-VE-T1A groups and A3KO-VE or A3KO-VE-B1A groups. **(D**,**E)** Photomicrographs showing expression of VE-T1A (**D)** and VE-B1A **(E)** in the NAc as revealed by immunostaining. The sections were co-stained for FOXP1, a marker for MSNs. **(F**,**G)** Quantification of the VE-T1A and VE-B1A expression by western blot. **(F)** Representative western blot showing the AAV-driven expression of VE-T1A and VE-B1A. **(G)** Quantification of the Arr3 expression shown in **F** (N=7-8).

We found that Arr3-derived peptide T1A rescued the hyperlocomotion in response to both AMPH (**Fig. 7B, Fig. S4, video 1 and 2**) and COC (**Fig. 7C**) to the same extent as full-length Arr3. In contrast, B1A was ineffective. Both peptides were expressed in targeted area, as detected by immunohistochemistry (**Fig. 7D,E**). Western blots confirmed the expression of both T1A and B1A peptides in NAc **(Fig. 7F)**. The expression of VE-B1A was significantly higher than that of VE-T1A (**Fig. 7F,G**).

### Arr3-dependent JNK3 activation is essential for the locomotor effects of psychostimulants

To further test the role of Arr3-dependent JNK3 activation in the locomotor effects of psychostimulants we used a mutant of another Arr3-derived JNK-activating peptide, T16 (**Fig. S2A-C, Fig S5A-C**) (Perry-Hauser et al., 2022), T16-F10V (**Fig. S5A**), which suppresses JNK activation (**Fig. S5B**,**C**). Cre-dependent expression of VE-T16-F10V in NAc of WT mice significantly reduced the locomotor response to both AMPH (**Fig. 8A**) and COC (**Fig. 8B**).

**Figure 8.**
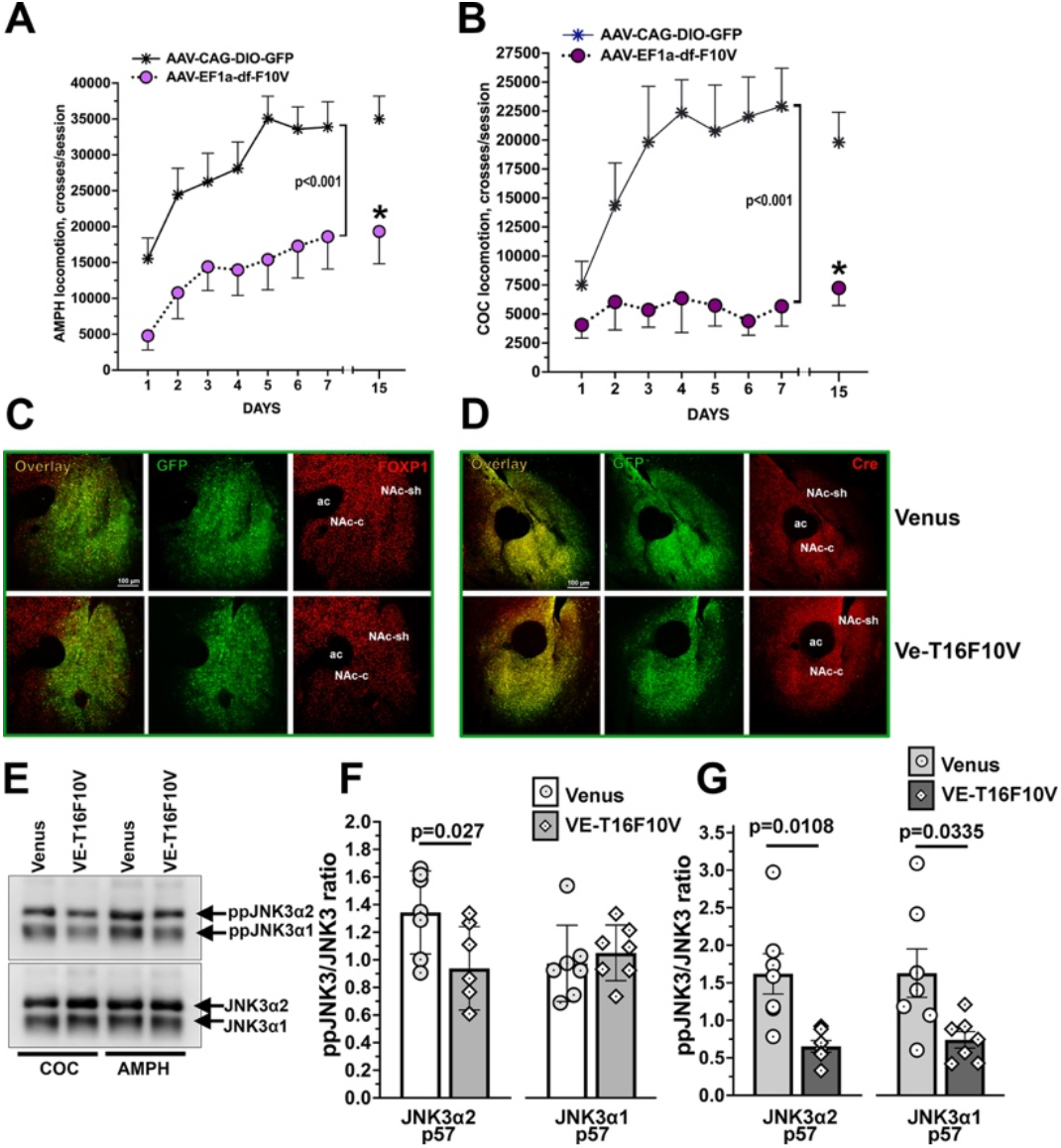
Arr3 mediates locomotor effects of psychostimulants by facilitating the activation of JNK3. **(A**,**B)** A3KO D1R-Cre mice were injected with Cre-dependent AAV encoding Venus or VE-T16-F10V and tested for drug-induced locomotion for 7 consecutive days and then for 1 day after a 7-day break. **(A)** AMPH-induced locomotion. **(B)** COC-induced locomotion. Bracket shows significant difference between the A3KO-VE and A3KO-VE-T16-F10V groups. (**C**,**D)** Immunohistochemical detection of Venus or VE-T16-F10V in the NAc. (**C**) The sections were stained with anti-GFP antibody and co-stained for the marker of MSNs striatal neurons FOXP1. **(D)** Adjacent sections through NAc stained for GFP to detect Venus or VE-T16F10V co-stained with anti-Cre antibody. **(E**,**F**,**G)** JNK3 was immunoprecipitated with anti-JNK3 antibody and blotted for doubly phosphorylated (active) JNK3 (ppJNK3). **(E)** Representative western blots showing immunoprecipitation. **(F)** Quantification of active JNK3 in the NAc following the AMPH treatment and testing (N=6-7). **(G)** Quantification of active ppJNK3 in NAc following the COC testing (N=7).

## Discussion

Mice lacking one of the two ubiquitous arrestin isoforms, Arr3, have been reported to be hyposensitive to the effect of the psychostimulant AMPH (Bouthenet et al., 1991; Beaulieu et al., 2005; Zurkovsky et al., 2017), demonstrated reduced locomotor responsiveness to morphine (Bohn et al., 2003; Urs et al., 2011), and blunted locomotor sensitization (Zurkovsky et al., 2017). Unexpectedly, the loss of Arr3 was reported not to affect the locomotor response to another psychostimulant, COC (Bohn et al., 2003; Gainetdinov et al., 2004). We found that A3KO mice are deficient in the locomotor responsiveness and sensitization to both psychostimulant drugs (**Figs. 1, 6**). These data suggest that Arr3 plays a role in the common neuronal mechanism involved in the psychostimulant-induced hyperlocomotion and locomotor sensitization.

Both AMPH and COC indirectly enhance striatal dopaminergic neurotransmission. Elevated dopamine (DA) in the main dopaminoreceptive areas results in excessive stimulation of dopamine receptors, primarily the D1R and D2R, and consequent hyperlocomotion [reviewed in (Baik, 2013; Johnson and Lovinger 2016)]. The DA receptors are G protein-coupled receptors (GPCRs) subject to homologous desensitization, which includes two steps: receptor phosphorylation by a G protein-coupled receptor kinase followed by arrestin binding to activated phosphorylated receptor. This stops (arrests) further G protein activation (Carman and Benovic, 1998). However, if the main Arr3 action were receptor desensitization, A3KO mice would be expected to show gain-of-function phenotype. For example, A3KO mice demonstrate enhanced morphine analgesia and reward (Bohn et al., 1999; Bohn et al., 2003), which is consistent with impaired desensitization of opioid receptors resulting in their overactivity. In the same vein, mice lacking another arrestin isoform, arrestin-2, showed enhanced locomotor responsiveness to AMPH (Zurkovsky et al., 2017). However, the phenotype in A3KO mice is loss-of-function, with sensitization to both AMPH and COC significantly suppressed. This suggests participation of Arr3-mediated signaling.

The CPu has been shown to play the leading role in in the psychostimulant-induced hyperlocomotion and sensitization (Kravitz et al., 2010; Ferguson et al., 2011; Kreitzer and Berke, 2011). Therefore, we expected that restoration of Arr3 to the CPu of A3KO mice would rescue their locomotor responses. Unexpectedly, exogenous Arr3 further inhibited already reduced locomotion in these mice (**Fig. 1**). Importantly, Arr3 expressed in the CPu of WT mice had the same effect (**Fig. 2**). The most parsimonious explanation of these findings is that in the CPu Arr3 acts as a negative regulator of the DA receptor signaling, thereby reducing the behavioral effects of psychostimulants. We found that the effect of Arr3 is driven by its activity in the D1R-bearing medium spiny neurons (MSNs) of the CPu: its selective expression in these neurons had a very strong locomotion-suppressing effect, whereas the expression in the D2R-bearing MSNs was ineffective (**Fig. 2**). This is consistent with previous findings that activation of D1R-bearing, but not D2R-bearing, MSNs enhances spontaneous (Kravitz et al., 2010) and AMPH-induced locomotion (Ferguson et al., 2011), as well as locomotor sensitization to AMPH (Ferguson et al., 2011).

Thus, the changes in the CPu do not explain the A3KO phenotype. However, the restoration of Arr3 to the NAc did rescue psychostimulant-induced hyperlocomotion and sensitization in A3KO mice **(Figs. 3-6)**. Thus, the phenotype of A3KO mice is driven by the loss of Arr3 in NAc, not in CPu. We probed the neuronal subtypes involved in the Arr3 action via cell-specific expression. We found that the expression of Arr3 in D1R-but not D2R-bearing neurons fully rescued hyperlocomotion in response to both AMPH and COC (**Figs.3-6)**. This is consistent with earlier reports of the leading role of the D1R-bearing MSNs in NAc in locomotor sensitization to psychostimulants (Hikida et al., 2010; Lobo et al., 2010) and in long-term drug-induced plasticity leading to behavioral changes in general (Lee et al., 2006; Grueter et al., 2013; MacAskill et al., 2014; Baimel et al., 2019).

It is important to note that in the dorsal striatum (CPu) the D1R-bearing MSNs give rise to the direct pathway projecting to the substantia nigra reticulata/globus pallidus internal, whereas the D2R-bearing MSNs give rise to the indirect pathway via projections to the globus pallisus external (Gerfen and Surmeier, 2011; Gerfen, 2022). Such discreet separation does not exist in NAc: although the D2R-bearing MSNs project to the ventral pallidum, the indirect pathway output structure of the ventral striatum, the D1R-bearing MSNs project to both the ventral striatum and the ventral tegmental area/substantia nigra, the output structures of both pathways (Smith et al., 2013; Kupchik et al., 2015). Although earlier studies suggested that striatal or accumbal D1Rs control acute responses to psychostimulants but have limited effect on locomotor sensitization (Vanderschuren and Kalivas, 2000; Baik, 2013; Johnson and Lovinger 2016), recent studies using optogenetic or DREADD-based inactivation of D1R-bearing MSNs demonstrated the key role of these neurons in locomotor sensitization to psychostimulants (Hikida et al., 2010; Ferguson et al., 2011; Lobo and Nestler, 2011; Chandra et al., 2013). D2Rs appear to play minimal role in locomotor responses to COC: COC-induced hyperactivity and locomotor sensitization were unchanged in D2R knockout mice (Sim et al., 2013). However, the role of D2R-bearing MSNs in CPu and NAc is poorly defined (Ferguson et al., 2011; Chandra et al., 2013; Gore and Zweifel, 2013; Rose et al., 2018; Donthamsetti et al., 2020).

We showed that Arr3 acts in the D1R-bearing MSNs, but it remains to be elucidated whether these D1R-bearing neurons belong to the direct or indirect pathway, or both. The relative role of Arr3 in D1R- and D2R-bearing MSNs in the control of locomotor responses to psychostimulants and locomotor sensitization was tested previously using cell type-specific Arr3 deletions and G protein- or arrestin-biased dopamine receptor mutants and ligands, but the results are contradictory (Allen et al., 2011; Urs et al., 2011; Peterson et al., 2015; Urs et al., 2016; Rose et al., 2018; Porter-Stransky et al., 2019; Donthamsetti et al., 2020). Some of the controversy, particularly in the studies involving biased signaling, might arise from the dual effects that the manipulations of Arr3 have on G protein-mediated versus Arr3-mediated signaling and, consequently, locomotor response. Interestingly, the results of rescue using DYN and ENK promoters did not fully agree with the outcome of Cre-driven neuron-specific expression. Arr3 expression in D1R-bearing MSNs via DYN promoter or D1R-Cre fully rescued the AMPH-induced locomotion. However, while A2A-Cre-dependent Arr3 expression was completely ineffective, Arr3 under ENK promoter yielded partial rescue (**Figs.4-5)**. We have previously shown that these promoters are selective for the DYN- and ENK-expressing neurons, respectively (Ahmed et al., 2024), in agreement with the previous report (Ferguson et al., 2011). In the CPu of hemiparkinsonian A3KO mice DYN-driven expression of Arr3 afforded full rescue of behavioral sensitization to L-DOPA, whereas ENK-controlled expression failed to do so (Ahmed et al., 2024). Conceivably, co-localization of DYN/D1R and ENK/D2R in NAc is not as strict as in CPu. Some D1R-expressing neurons, particularly those residing the NAc shell giving rise to the indirect pathway and projecting to the dorsolateral ventral pallidum (Zhou et al., 2003) co-express ENK. There is ample evidence that DYN and ENK in CPu hardly overlap (Lu et al., 1998; Zhou et al., 2003), but little is known about the degree of segregation of the DA receptors and neuropeptides in NAc.

Enhanced responsiveness to direct or indirect GPCR activators in mice lacking arrestins is easily explained by reduced receptor desensitization and consequent prolonged G protein-mediated signaling. In contrast, loss-of-function phenotype suggests that arrestin-mediated signaling plays a role: when it is missing due to absence of Arr3, the behavioral responsiveness is diminished. This suggests that in psychostimulant-induced locomotor responses Arr3 acts as a signal transducer downstream of the dopaminergic signaling in the D1R-bearing neurons in NAc, presumably, via D1R. We showed that Arr3 supports psychostimulant-induced hyperlocomotion and locomotor sensitization by facilitating JNK3 activation. This is evidenced by the fact that JNK3-activating T1A (Zhan et al., 2016; Perry-Hauser et al., 2022), a short 25-residue peptide that lacks receptor-binding elements and does not perform other signaling functions of Arr3, rescued psychostimulant-induced hyperlocomotion of A3KO mice to the same extent as full-length Arr3 (**Fig. 7**). Furthermore, the expression in NAc of a mutant Arr3-derived peptide T16-F10V that inhibits JNK activation markedly reduced the activation of JNK3 and the responsiveness of WT mice to AMPH (**Fig. 8**). Thus, Arr3 supports psychostimulant-induced hyperlocomotion by facilitating JNK3 activation.

Sensitization defined as augmentation of behavior upon repeated treatment with a drug is known to be caused by drugs of different types engaging directly or indirectly with the DA transmission in the core dopaminoreceptive regions such as CPu and NAc (Schmidt and Beninger, 2006; Delage et al., 2023). This includes sensitization of rotational behavior and of abnormal involuntary movements (AIMs) in hemiparkinsonian rodents induced by DA precursor L-DOPA or DA agonists (Schmidt and Beninger, 2006; Abe et al., 2023; Ahmed et al., 2024) and locomotor sensitization to psychostimulants such as AMPH and COC (Schmidt and Beninger, 2006; Zurkovsky et al., 2017). It stands to reason that these sensitization phenomena have mechanistic commonalities. In fact, some common neural and molecular alterations associated with the sensitization process have been noted (Nestler et al., 1999; Graybiel et al., 2000; Beck et al., 2019; Savell and Hope, 2023; Zamanian et al., 2025). We previously found that Arr3 mediates abnormal locomotor responses to L-DOPA in hemiparkinsonian mice via facilitation of JNK3 activation (Ahmed et al., 2024). Combined with the data presented here this suggests that Arr3-dependend regulation of the JNK3 activity is a common molecular mechanism playing an important role in the sensitization process caused by different drugs. Furthermore, in both cases Arr3 in D1R-bearing neurons is specifically involved, albeit in different striatal subdivisions.

## Conclusions

Arr3 supports the locomotor response to psychostimulants via enhanced activation of JNK3 in the D1R-bearing neurons in NAc. Thus, in these neurons Arr3 performs a signaling function critical for the responsiveness and locomotor sensitization. The evidence presented here, together with our previous results regarding the locomotor responsiveness to dopaminergic stimulation by L-DOPA (Ahmed et al., 2024), suggest that Arr3-assisted JNK3 activation is a previously unappreciated universal pathway mediating DA signaling in the striatum. Importantly, there is ample evidence that Arr3 regulates JNKs without recruitment to GPCRs (reviewed in (Gurevich and Gurevich, 2024)). The molecular mechanism of signal integration between activation of DA receptors following psychostimulant administration and Arr3 action on the JNK pathway remains to be elucidated.

## Acknowledgements

We thank Dr. Robert J. Lefkowitz (Duke University) for the gift of the arrestin-3 knockout mice. We are grateful to Dr. John F. Neumaier (University of Washington) for the gift of the rat enkephalin and dynorphin promoter vectors. This work was supported by NIH grants RO1 NS065868 and R21 DA030103 (to EVG) and R35 GM122491 (to VVG).

## Author contributions

MRA, JLD, CZ, SK, SM, CB, JJ-M, KY, HL and EGV conducted the study. MRA, JLD, CZ, EVG, and VVG designed the experiments. MRA, EVG and EVG wrote the paper.

## Methods

### Animals and tissue preparation

All animal procedures followed the guidelines in the Guide for the Care and Use of Laboratory Animals of the National Institutes of Health. The protocol was approved by the Institutional Animal Care and Use Committee of Vanderbilt University. A3KO mice were kindly provided by Dr. R. J. Lefkowitz (Duke University). Mice expressing Cre recombinase under control of D1R promoter (D1RCre) were purchased from the Jackson Laboratory (B6.FVB(129S6)-Tg(Drd1a-cre)AGsc/KndlJ strain on C57Bl/6J-congenic background). Mice expressing Cre recombinase under control of A2A adenosine receptor (A2ARCre) were kindly provided by Dr. Brad Grueter,

Vanderbilt University. All animals were housed at the Vanderbilt University animal facility with a 12/12 h light/dark cycle and free access to food and water. Mice were bred using heterozygous breeding pairs to obtain A3KO and wild type (WT) littermates. To maintain genetic homogeneity, all mice were consistently backcrossed to WT C57Bl mice purchased from Charles River. To minimize background genetic differences between the Cre lines, the same WT mice were used for backcrosses in both lines. To produce A3KO mice that also expressed Cre under neuron-specific promoters, Arr3+/- mice were bred with hemizygous Cre mice to produce A3KO/A1RCre+/- or A3KO/A2ACre+/- mice as well as their WT littermates that were used in the experiments. In cases when more A3KO mice than WT mice were needed, limited breeding of A3KO mice to Cre lines was employed to minimize the total number of used animals.

### Drug Treatment and Locomotor activity Measurements

Locomotor activity was measured in open field chambers (Med Associates Inc., Fairfax, VT) equipped with Activity software, as described (Zurkovsky et al., 2017). The data were collected in 5-min intervals (bins). The activity was first measured for 30 min before the injection of AMPH or COC. Following pre-drug testing, the mice received 3 mg/kg of AMPH or 20 mg/kg COC i.p. and were immediately placed back into the apparatus and tested for AMPH- or COC-induced locomotor activity for 90 min. The testing was performed daily for 7 days. After that, the mice were withdrawn from the drug for 7 days and re-tested again with the same drug dose on Day 15.

### Virus construction and preparation

The full-length coding sequence of the bovine Arr3 (Arr3) (gi 6978467) (a.k.a. β -arrestin2) was C-terminally tagged with HA (Tyr-Pro-Tyr-Asp-Val-Pro-Asp-Tyr-Ala). The viral construct assembled in the lentiviral vector (LV) pLenti6.4/V5-DEST included GFP under control of the CMV promoter and downstream co-cistronic Arr3-HA under control of super-IRES (sIRES), which has been shown to significantly increase protein expression (Bochkov and Palmenberg, 2006). LVs expressing GFP alone were used as a control. The rat enkephalin (ENK) and dynorphin (DYN) promoter vectors were a gift from Dr. Neumaier (the University of Washington) (Ferguson et al., 2011). The promoter sequences were inserted into promoterless pLenti6.4/V5-DEST vector. Venus-tagged Arr3 was inserted after sIRES. The lentiviruses were produced using the ViraPower system (Invitrogen, Carlsbad, CA), concentrated and purified as described (Ahmed et al., 2010). Viral titers of LVs with the CMV promoter were measured based on GFP or HA expression using HEK293 cells infected with the appropriate viruses. The titers of LVs with cell-specific promoters were determined by Western blot for HIV1 p24 band using mouse monoclonal antibody (ThermoFisher).

The adeno associated viruses serotype 5 (AAV5) for pan-neuronal expression under control of CMV promoter encoding Venus (control), VE-Arr3, VE-T1A, VE-B1A, or VE-T16-F10V was constructed using the pAAV-MSC vector from the AAV-5 Promoterless Helper Free Expression System and 293-AAV cell line (Cell Biolabs, Inc.).

AAV clones for neuron-specific Cre-dependent expression under control of the EF1α promoter were constructed using a double-floxed inverted open reading frame backbone (AAV p-EF1α-DIO-WPRE-hGHpA from Addgene) to ensure Cre-dependent expression and produced using AAV-5 Promoterless Helper Free Expression System and 293-AAV cell line (Cell Biolabs, Inc.).

The viral titer was determined by standard qPCR with the BioRad SYBR Green supermix using serial dilutions of the cloning vector pAAV-MCS as standards.

### Animal surgeries and virus injection

The LVs or AAVs containing Arr3 (or GFP control) were injected bilaterally into the dorso-lateral posterior CPu, at coordinates AP +1.02; ML 1.65; DV 3.55. Alternatively, the injections were made bilaterally into the NAc, at coordinates AP +1.60; ML ±0.62 (bilateral); DV 4.70. The mice were anesthetized with ketamine/xylazine (100/10 mg/kg i.p.) and mounted on an automated digital stereotaxis (Kopf Instruments). The virus injection (3 μl of the concentrated virus in saline per CPu and 1μl per NAc 0.3 μl/min) was made with an automatic injector, and the needle was left in place for another 10 min. The skin of the scull was then sutured, and the mice were allowed to recover from anesthesia on a heated pad before return to their home cages. The behavioral experiments started 4 weeks after the virus injection.

### Tissue preparation

Upon completion of the drug administration and behavioral testing, the mice were euthanized with isoflurane, decapitated, the brains were collected and rapidly frozen on dry ice. The rostral parts of the brains containing the striata were kept at -80^°^C until samples were collected for Western blot analysis. To collect the samples for Western blot, the brains were cut through the striatum, the position of the virus injection track was identified, and the tissue around the injection track was collected on both sides and pooled to be used to determine the expression of virus-encoded proteins, as described (Ahmed et al., 2010; Ahmed et al., 2015). Randomly selected animals were overdosed with ketamine/xylazine and transcardially perfused with 4% paraformaldehyde. The brains were removed, post-fixed in 4% paraformaldehyde, cryoprotected in 30% sucrose, kept frozen at -80^°^C, then cut into 30μM sections and used for immunohistochemistry

### Immunoprecipitation

Immunoprecipitation (IP) of JNK3 was performed as previously reported (Ahmed et al., 2024). The nucleus accumbens from control, AMPH and COC treated mice were dissected on ice and rapid frozen using liquid nitrogen until further processing and stored at -80°C. The tissue samples were lysed with lysis buffer containing protease and phosphatase inhibitors and sonicated on ice to minimize damage and processed for IP. The lysates were rocked for 30 mins at 4°C and centrifuged to isolate the supernatant. The supernatant was then precleared with protein G plus agarose (Santa Cruz Biotechnology, CA). 2mg of the protein lysate was used to IP JNK3 using JNK3 specific rabbit monoclonal antibody (Cell Signaling cat#2305; 1:100) for 2hours at 4°C. Protein G plus agarose slurry was then added to the mixture to trap antibody bound JNK3 to the agarose beads overnight at 4°C. The agarose beads were then carefully isolated by centrifugation and transferred to fresh durapore PVDF centrifugal filters (Millipore, Cat # UFC30DV00, CA). The beads were repeatedly washed with lysis buffer, and the samples were eluted by heating to 90-95°C in a heat block using 100µl of 2X Laemmli sample buffer with β-mercaptoethanol added to the buffer. The samples were blotted for total JNK3 and phospho JNK antibodies (Cell Signaling, cat#2305 and 4668 respectively). The values were measured in arbitrary units in terms of active JNK over total JNK3 that was immunoprecipitated.

### Western blot

The expression of GFP- or Venus-tagged constructs was measured by Western blot with mouse anti-GFP antibody (Clontech; 1:2,000). The expression of WT and mutant Arr3 was measured with rabbit polyclonal anti-Arr3 antibody, as described (Ahmed et al., 2007). To detect HA-tagged constructs, rabbit monoclonal anti-HA antibody (Cell Signaling, cat.# 3724; 1:1,000) was used. Anti-rabbit or anti-mouse horseradish peroxidase-conjugated secondary antibodies (Jackson Immunoresearch) were used at 1:10,000 dilution. The blots were developed with a chemiluminescent substrate and either exposed to X-ray film or subjected to direct detection using C-DiGit Blot scanner (Li-Cor). The gray values of the bands on X-ray film were measured with a Versadoc imaging system (Bio-Rad) with QuantityOne software.

### Immunohistochemistry

To detect the expression of Venus-tagged constructs by immunohistochemistry, the mouse monoclonal anti-GFP (JL-8, Clontech; 1:500 overnight at 4^°^C) primary antibody was used, followed by goat anti-rabbit biotinylated secondary antibody and streptavidin conjugated with Alexa Fluor 488 (Invitrogen, Carlsbad, CA). For double labeling with FOXP1, the sections were labeled with mouse anti-GFP (1:500) antibody and rabbit anti-FOXP1 antibody (Cell Signaling Technology; 1:400) and detected by biotinylated secondary antibody/streptavidin-Alexa Fluor 488 (GFP; green) and anti-rabbit-Alexa Fluor 568 (FOXP1; red). Alternatively, chicken anti-GFP antibody (Invitrogen; 1:500) was used to label Venus-tagged Arr3. For double-labeling with Cre, the sections were labeled with chicken anti-GFP (1:500) antibody and rabbit anti-Cre antibody (Cell Signaling Technology; 1:400) and detected by biotinylated secondary antibody/streptavidin-Alexa Fluor 488 (GFP; green) and anti-rabbit-Alexa Fluor 568 (Cre; red). The sections were photographed at low magnification on a Nikon TE2000-E automated microscope with a 4x dry objective and an Andor Zyla high resolution digital camera using the stitching function of the Nikon NIS-Elements software. High power photographs were collected on an Olympus FV-100 confocal microscope in the green and red channels with z-sectioning using a 40x oil immersion objective at 1024 x 1024 pixels. The images were assembled in Photoshop, with minimal contrast adjustments applied separately to channels to equalize their intensity.

### Quantification and statistical analysis

The locomotion data were analyzed by two-way repeated measure ANOVA with Group (WT-GFP versus A3KO-VE, A3KO-Arr3) as a between group and Day as a repeated measure factor. The group differences across sessions were assessed by the post hoc Bonferroni/Dunn, Tukey, or by Dunnett’s test (with correction for multiple comparisons) where appropriate. When a significant effect of Group was observed, the data for individual days were compared by two-way repeated measure ANOVA with Group (WT-GFP versus A3KO-VE, A3KO-Arr3) as a between group and Bin as a repeated measure factor followed by the Bonferroni post hoc test, when appropriate. The molecular data were analized by one-way ANOVA with Dunnett’s post hoc comparison or by unpaired t-test with Welsh’s correction where appropriate.

Statistical analysis of the data was performed using Prizm 10. In all cases, p<0.05 was considered significant.

