## Supplemental Data for "Arrestin-3 promotes locomotor sensitization to psychostimulants via JNK signaling in nucleus accumbens"

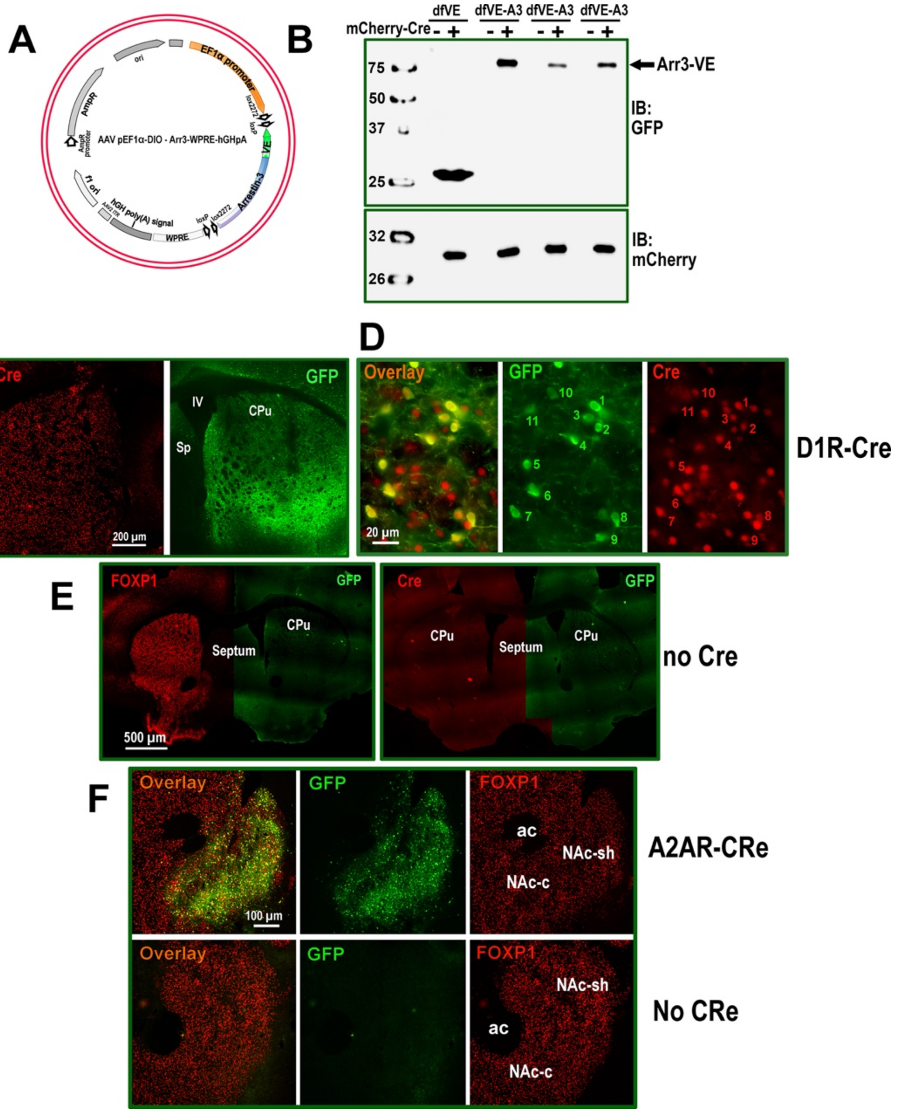


**Figure S1. The AAV clones and Cre-dependent expression. (A)** The design of the AAV clone with double-floxed open reading-frame inverted arrestin-3 tagged on N-terminus with Venus. The expression of arrestin-3 is driven by EF1$\alpha$ promotor ensuring a modest level of expression. **(B)** The experiment showing strict Cre-dependence of arrestin-3 expression driven by the AAV-p EF1α-DIO-dfVe-Arr3 shown above. HEK293 cells were transfected with the AAV vector and co-transfected with mCherry-tagged Cre or empty vector. The Western blot analysis demonstrated the VE-Arr3 expression only in cells co-expressing Cre. **(C)** The expression of Venus-tagged arrestin-3 in CPu of D1RCre mice. The sections were co-stained for GFP, to detect arrestin-3, and for Cre. **(D)** High power photomicropgraphs showing an overlap of Cre with VE-Arr3 (see corresponding numbers). **(E)** The lack of VE-Arr3 expression in a Cre-negative mouse. The sections were co-stained for a marker of striatal neurons FOXP1 (left panel) or Cre (right panel). **(F)** the expression of VE-Arr3 in the NAc of A2Acre mouse (upper panel) and the lack of VE-Arr3 expression in a Cre-negative mouse (lower panel)

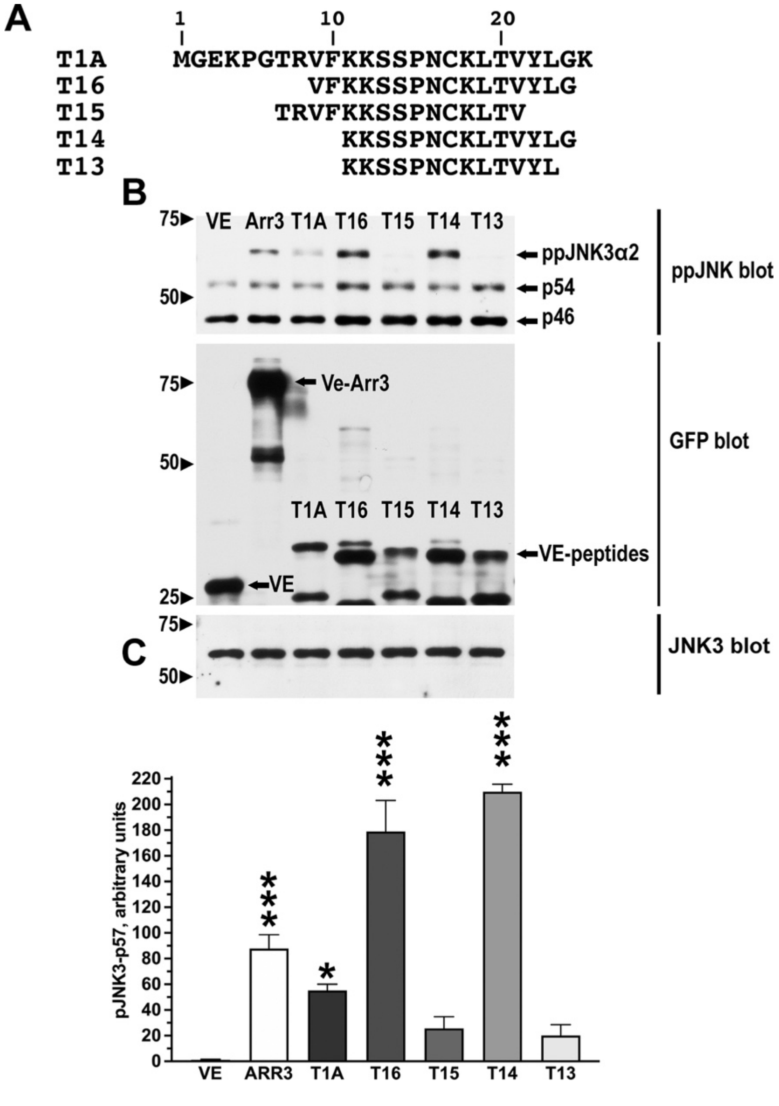


**Figure S2. The arrestin-3-derived peptides activate JNK3.** Neuro2A cells were transfected with HA-JNK3α2, Venus control, full length arrestin-3, or indicated Venus-tagged arrestin-3-derived N-terminal peptides. The JNK activation was detected with anti-phospho-JNK antibody; the expression of arrestin-3 and the peptides – with anti-GFP antibody. **(A)** Structure of the arrestin-3-derived N-terminal peptides. **(B)** Representative Western blots showing the JNK activation and peptide and JNK3 expression. **(C)** Quantification of the JNK3$\alpha$2 phosphorylation induced by the peptides. The amount of ppJNK3$\alpha$2 was normalized to the amount of the total JNK3 in the same samples.

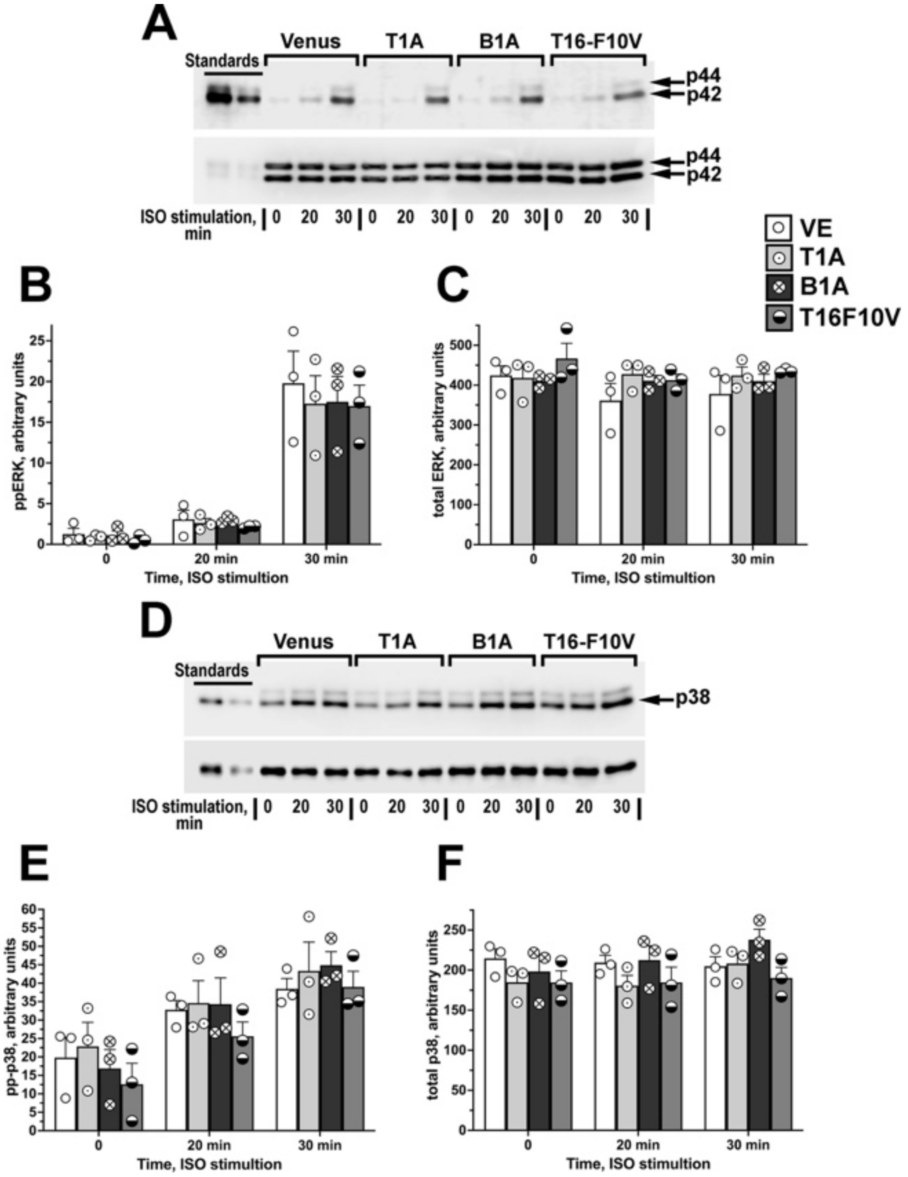


**Figure S3. Arrestin-3-derived peptides do not affect the activity of ERK or p38. (A)** Representative Western blot of the ERK1/2 activation following stimulation of HEK DKO cells with isoproterenol (ISO, 10 $\mu$M) for indicated time with or without co-expression of peptides. Venus was used as a negative control. Upper panel – double phosphorylated (active) ERK1/2; lower panel – total ERK1/2 expression. **(B,C)** Quantification of the Western blot data for phospho-ERK **(B)** and total ERK **(C)** (N= 3). **(D)** Representative Western blot of the p38 activation following stimulation of HEK DKO cells with isoproterenol (ISO, 10 μM) for indicated time with or without co-expression of peptides. **(E,F)** Quantification of the Western blot data for phospho-p38 **(E)** and total p38 **(F)** (N= 3).

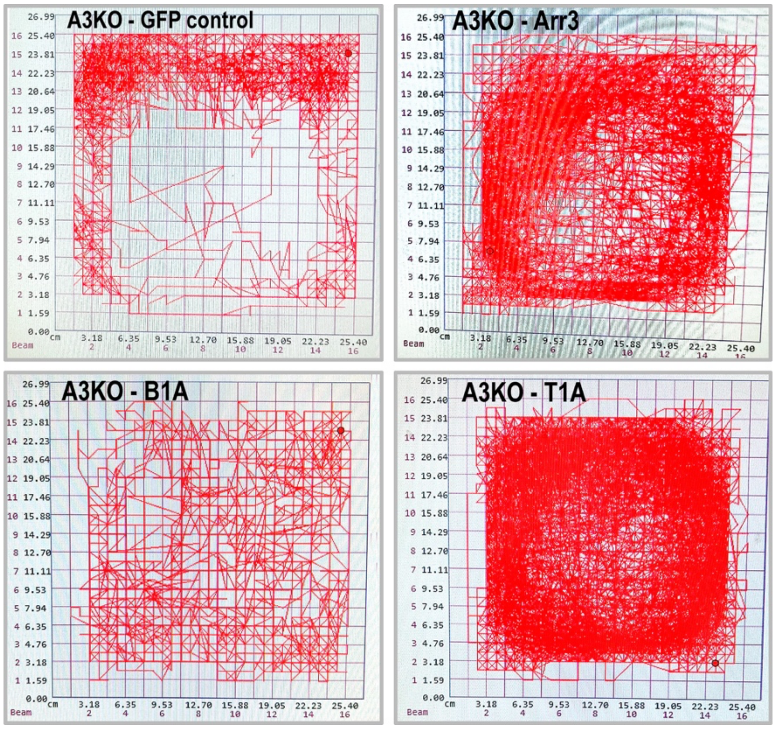


**Figure S4. The arrestin-3-derived peptide T1A, but not arrestin-2-derived peptide B1A, promotes amphetamine-induced hyperlocomotion.** Screenshots of automatic tracing of the amphetamine-induced hyperlocomotion in A3KO mice injected into the nucleus accumbens with indicated constructs.

**Video 1.** Sample recording of the COC-induced locomotion in A3KO mouse expressing wild type VE-Arr3 in the NAc.

**Video 2.** Sample recording of the COC-induced locomotion in A3KO mouse expressing Arr3-derived peptide VE-T1A in the NAc.

**Video 3.** Sample recording of the COC-induced locomotion in A3KO mouse expressing Arr2-derived control peptide VE-B1A in the NAc.

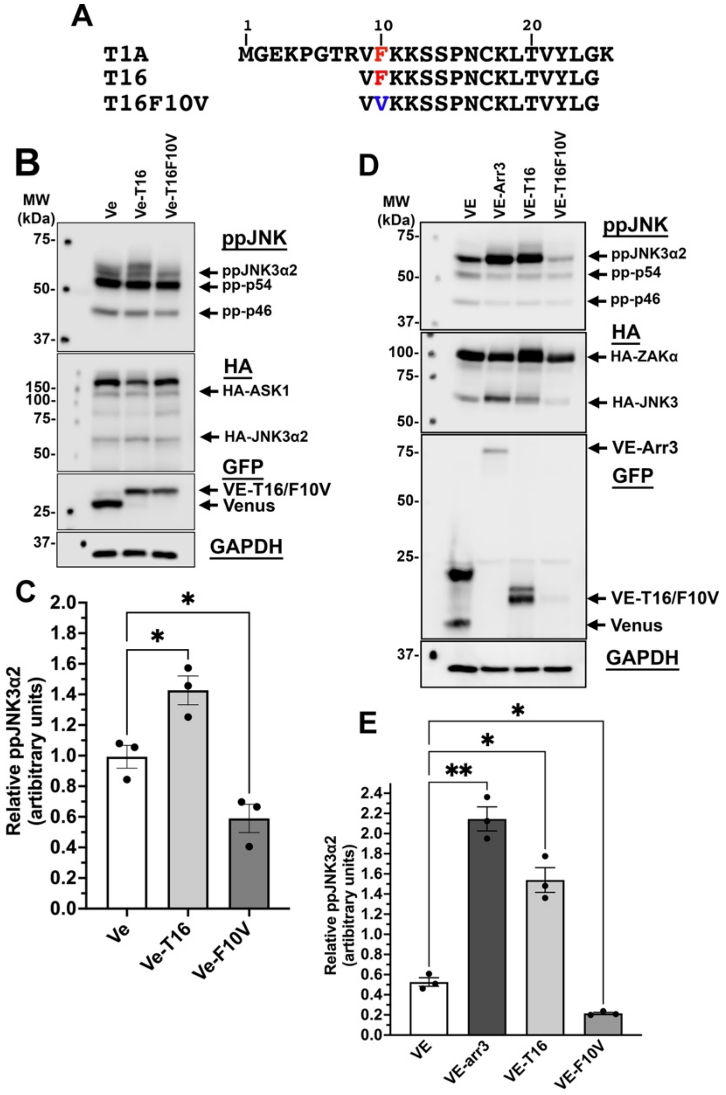


**Figure S5. Mutant T16F10V inhibits the activity of JNK3. (A)** Structure of T1A, T16 and mutant T16-F10V. **(B)** A representative Western blot showing that T16F10-V inhibits ASK1-promoted activation of JNK3 in HEK293 AKO cells. **(C)** Quantification of the Western blot data (N=3). (D) A representative Western blot showing that T16F10-V inhibits ZAKα-promoted activation of JNK3 in HEK293 AKO cells. The statistical analysis was performed by one-way ANOVA followed by Dunnett's post hoc comparison. *, $p<0.05$
